# Playback calls help to increase the detectability of *Coturnix coturnix* (Common quail), a cryptic and widespread galliform

**DOI:** 10.64898/2026.02.13.705393

**Authors:** Eduardo Laguna, Inmaculada Navarro, Raquel Castillo-Contreras, José A. Torres, Juan Rubiales, Miguel Beloki, Carlos Sánchez-García

**Author notes:** Correspondence author: Inmaculada Navarro, Fundación Artemisan, 13001, Ciudad Real, Spain.

## Abstract

In cryptic or difficult-to-detect bird species, the monitoring schemes based on generalist detection methods may introduce bias into abundance estimates and population indices. This the case of the *Coturnix coturnix* (Common quail), a migratory Palearctic galliform, in which the use of passive detection methods within breeding birds monitoring schemes may not be efficient owing to its complex socio-sexual system and migratory behavior.

For the first time, *C.coturnix* detectability was simultaneously compared using standard passive, generalist multispecies survey methods from the Pan-European Common Bird Monitoring Scheme (PECBMS) and a species-specific active survey employing female call playback. Surveys were conducted at 1,077 listening points within 107 transects over four breeding seasons (2022–2025) in open farmland landscapes dominated by cereal crops in Extremadura, south-western Spain. Detection counts differed substantially between methods: active surveys increased expected counts by 72% (95% CI: 59–85%) compared to passive surveys. The increase in *C.coturnix* detections elicited by playback showed a non-linear, density-dependent pattern, being highest at low passive abundances per listening point (maximum at 3–4 individuals) and stabilizing at intermediate abundances. This indicates that call playback is particularly effective at detecting individuals that would otherwise remain undetected. Our findings suggest that passive, multispecies surveys may underestimate *C.coturnix* abundance, especially in low-density populations. Integrating species-specific active methods into monitoring programs can improve detectability, generate more reliable population indices, and support evidence-based conservation and management strategies for this elusive species.

**LAY SUMMARY:**

- Bird monitoring schemes guide conservation decisions across Europe, but generalist schemes based on passive methods may miss species that are hard to detect such as *C.coturnix* Common quail, a migratory farmland bird that hides in dense crops. In practice, only males spontaneously calling can be detected, hence passive methods could lead to underestimates of its abundance and even false absences in low-density areas.
- We compared standard passive surveys with surveys that added a recorded female call (playback) to stimulate male responses. Across 1,077 listening points monitored over four breeding seasons in southwestern Spain, playback increased the number of birds detected by 72% compared with passive methods alone.
- The improvement was strongest where *C.coturnix* numbers were low, showing that many individuals remain undetected without playback. Incorporating simple, species-specific methods into monitoring programs can produce more reliable population estimates and strengthen conservation and management decisions for this elusive species.

## INTRODUCTION

In Europe, the Pan-European Common Bird Monitoring Scheme (PECBMS) is the official breeding bird monitoring scheme (BMS), which aims to track changes in breeding populations of common birds and produce indices that reflect the overall state of nature (https://pecbms.info/about-us). As other BMS around the world, PECBMS provides essential long-term data on bird populations, supporting biodiversity assessments, conservation planning, and policy decisions (Gregory et al. 2005, Moussy et al. 2022). By integrating standardized protocols from national monitoring schemes, PECBMS generates trends and indices for over 170 species, informing conservation assessments and evaluating the impact of policies such as the European Union Common Agricultural Policy.

Despite their success, generalist BMS have limitations when applied to species whose behavior, spatial ecology, or social organization deviate from standard survey assumptions. Detection probability can vary across species, habitats, and behavioral states, introducing bias into abundance estimates and population indices (Nichols et al. 2007, Kéry and Schmidt 2008, Sardà-Palomera et al. 2012a). These challenges are particularly pronounced for cryptic, elusive, or patchily distributed species, where passive surveys may fail to capture true abundance, being this the case for *Coturnix coturnix* (Common quail).

This cryptic, terrestrial galliform is widespread across Eurasia, but its breeding biology, complex socio-sexual system, and migratory behavior make it difficult to detect. Most of its life cycle occurs within dense cereal crops, and males are primarily detectable through their breeding calls, while females and paired males often remain undetected (Guyomarc’h et al. 1998, Rodríguez-Teijeiro and Puigcerver 2020). Vocalizations serve multiple functions, including mate attraction, location signaling, and individual recognition (Collins and Goldsmith 1998, Derégnaucourt and Guyomarc’h 2003), and in males, calling activity depends on the presence and behavior of conspecifics (Pincemy and Guyomarc’h 2004). Consequently, passive survey methods that record only spontaneously calling males may substantially underestimate population size.

Previous studies comparing passive and active survey methods were conducted on consecutive days, with passive surveys performed first and active surveys on the following day (Puigcerver et al. 2018, Sardà-Palomera et al. 2022). Given the high mobility of *C*.*coturnix* during the breeding season (Munteanu and Maties 1974, Schleidt 1983, Rodríguez-Teijeiro et al. 1992, Sardà-Palomera et al. 2011), this sequential design could compromise the comparability of detection results.

To address these limitations, we conducted simultaneous passive and species-specific active surveys using female *C*.*coturnix* call playback across four breeding seasons in Extremadura, south-western Spain. Our objectives were: (i) to assess whether active playback calls improves quail detectability relative to passive methods, and (ii) to examine whether the relationship between passive and active detections is linear or density-dependent. These results aim to guide the development of robust monitoring protocols applicable across the species’ range.

## MATERIALS AND METHODS

### Study area

This study was conducted from 2022 to 2025 in 68 10×10 km UTM grid cells located in the region of Extremadura, south-western Spain (Figure 1). Study sites encompassed habitats suitable for quail, more specifically open farmland landscapes dominated by cereal crops.

**FIGURE 1.**
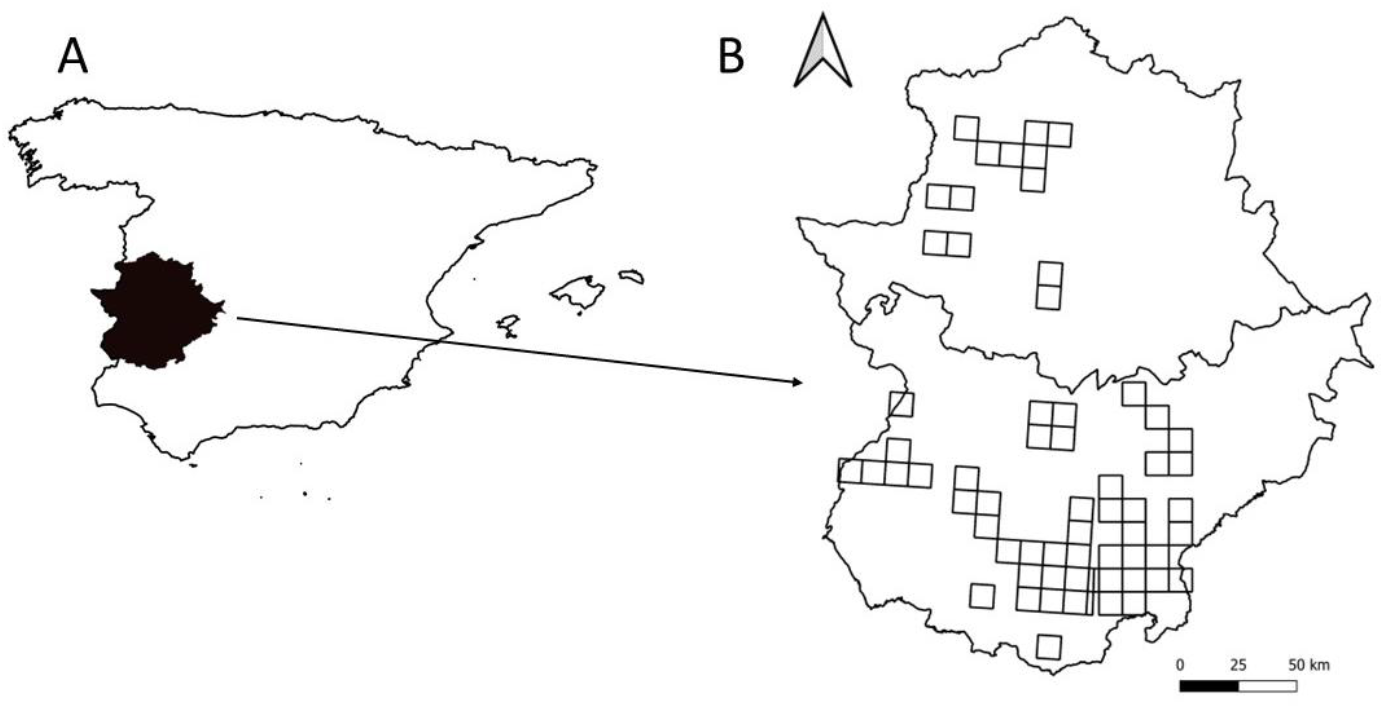
Study area (peninsular Spain and Balearic Islands context) (**A**) and location of the field sites within 68 10×10 km UTM grid cells in Extremadura (**B**).

**FIGURE 2.**
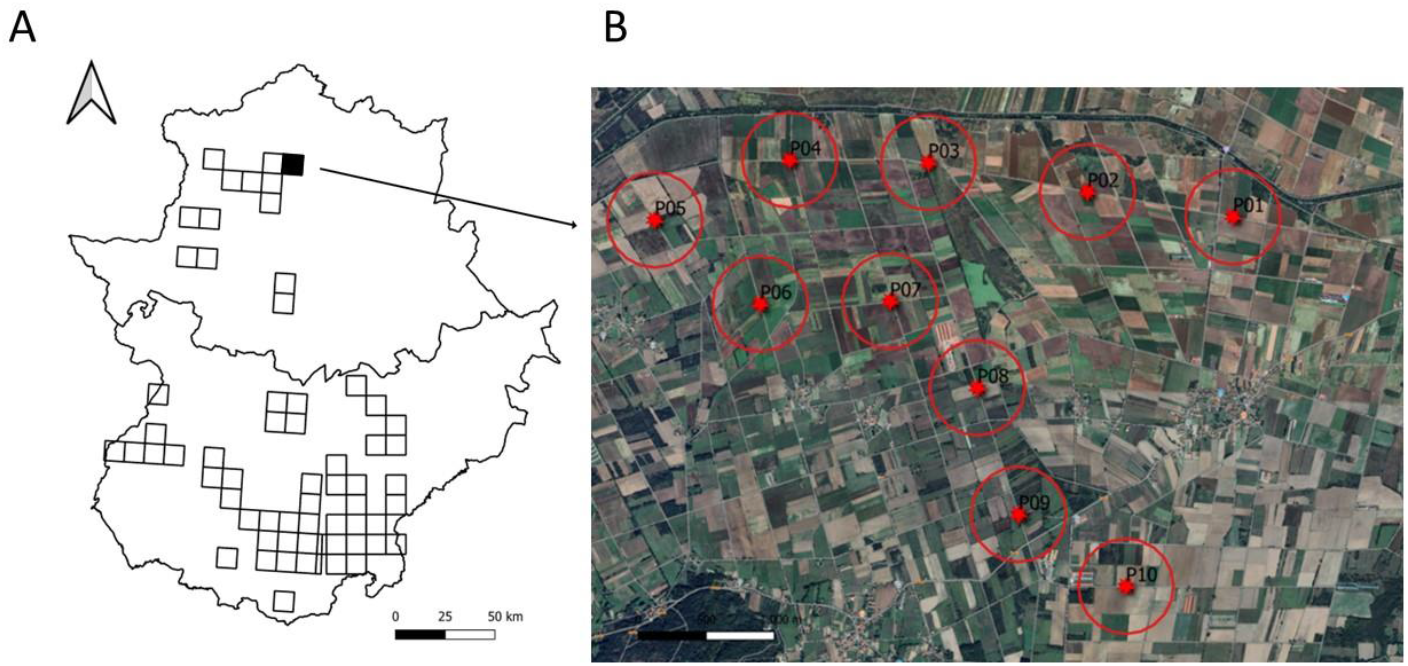
Location of 68 10×10 km UTM grid cells where *C*.*coturnix* surveys were conducted (**A**), and example of a survey transect with 10 listening points (**B**).

### Survey methods

Surveys were conducted during the *C*.*coturnix* breeding season (April-June) under suitable weather conditions, primarily during periods of expected peak vocal activity, including early morning and late afternoon hours. Each survey transect comprised 10 listening points spaced at least 700 m apart to avoid overlap in detection.

At each point, observers first conducted a 3-minute passive listening period (recording all detected *C*.*coturnix*), followed immediately by 1-minute playback of a female call and a subsequent 3-minute active listening period (recording all detected *C*.*coturnix*). This nested structure (listening points within surveys, surveys across years) implies non-independence among observations, which was explicitly accounted for in the statistical analyses. At each listening point, *C*.*coturnix* detected were recorded using a digital app for wildlife and game monitoring (CensData), and on a paper-based protocol (Supplementary Material Table S1).

The playback device consisted on a digital tape playing the female call to attract breeding males (Rodríguez-Teijeiro et al. 2019) and had an approximate action radius of 350 m (Sardà-Palomera et al. 2012b, Rodríguez-Teijeiro et al. 2019).

### Data analysis

Counts obtained without playback (N_passive) and with playback (N_active) were numeric variables. Year, UTM grid cell, and listening point identity were treated as categorical factors. To uniquely identify sampling locations, combining UTM grid cell and listening point number created a composite point identifier (ID_Point).

To allow direct comparison between survey methods within a single modelling framework, the dataset was reshaped to long format, creating a factor variable (Method) with two levels (passive vs active) and a corresponding response variable (N), representing the number of individuals detected per listening point and method.

### Hierarchical modelling of detection counts

Differences in *C*.*coturnix* detections between passive and active survey methods were analysed using a generalized linear mixed-effects model (GLMM) with a Poisson error distribution and log link function. The response variable was the number of detected individuals (N), and survey method (Method) was included as a fixed effect. The hierarchical generalized linear mixed model used to compare passive and active survey methods can be written as:

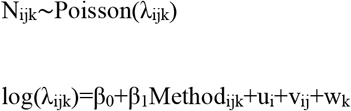

where N_ijk_ is the number of *C*.*coturnix* detected at listening point j within survey transect i during year k.

To account for the hierarchical structure of the data and repeated measurements, random intercepts were included for sampling points nested within survey transect (point/ survey transect) and for year:

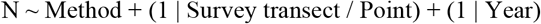

Model parameters were estimated using maximum likelihood with Laplace approximation, as implemented in the lme4 package (Bates et al. 2015) in R (R Core Team 2024). This hierarchical formulation explicitly accounts for the non-independence of repeated observations within sampling points and surveys, allowing inference on method-related differences in detection while controlling for spatial and temporal clustering.

### Model diagnostics and dependence assessment

Model adequacy and distributional assumptions were evaluated using simulation-based residual diagnostics implemented in the DHARMa package (Hartig and Hartig 2017). Uniformity, outliers, and dispersion of residuals were assessed through graphical inspection and formal tests.

A nonparametric dispersion test revealed significant subdispersion (dispersion = 0.31, *P* = 0.016), indicating that observations were more similar than expected under a purely Poisson process. This pattern is consistent with dependence among observations within sampling points and surveys, supporting the use of a hierarchical modelling framework.

The effect of survey method on detection rates was quantified using the fixed-effect estimate for the active method. Exponentiation of the model coefficient provided a multiplicative factor representing the expected increase in detected individuals when using call playback relative to passive surveys. Model-based predictions were generated on the response scale and compared with observed values to assess overall fit. Mean predicted detections and associated standard errors were calculated for each survey method.

### Density dependence in detection gain

To assess whether the gain in detections provided by call playback varied across density gradients, we explicitly modelled the increment in detections attributable to the active method. For each sampling point, we defined the detection increment as:

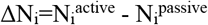

Where 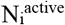 and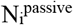 denote the number of *C*.*coturnix* detected during passive listening and subsequent call playback, respectively.

To evaluate potential non-linear density-dependent patterns in detection gain, we fitted a generalized additive model (GAM) with a Poisson error distribution and log link function:

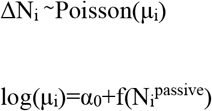

where f() is a smooth function describing how the increment in detections varies as a function of passive detections.

To account for repeated surveys at the same locations and interannual variability, random intercepts for sampling point and year were included in the model. This approach allowed us to identify non-linear density-dependent changes in the effectiveness of call playback, such as saturation effects at higher densities, while explicitly reflecting the sequential survey design (passive detection followed by active playback).

## RESULTS

Between April and June from 2022 to 2025, 1,197 listening points nested within 119 survey transects were conducted in Extremadura, sampling 68 different 10 × 10 km UTM grid cells in areas dominated by open agricultural habitats (Table 1 and Figure 1). However, as the aim of the study was to evaluate the difference in *C*.*coturnix* detection between active and passive methods, we excluded information from all survey transects in which 0 detections were obtained for both methods at all listening points. Therefore, a total of 2,154 observations from 1,077 sampling points nested within 107 surveys from four years were included in the hierarchical analysis.

**TABLE 1.** Summary table showing the number of sampled grids, survey transect, listening points, and *C*.*coturnix* detections during the spontaneous listening period (N_passive) and after playback (N_active) for the four years under study.

| Year | UTM Grid | Survey transects | Listening points | *C.coturnix* counts (N_passive) | *C.coturnix* counts (N_active) |
| --- | --- | --- | --- | --- | --- |
| 2022 | 15 | 15 | 157 | 258 | 414 |
| 2023 | 13 | 14 | 140 | 41 | 147 |
| 2024 | 36 | 36 | 360 | 243 | 380 |
| 2025 | 53 | 54 | 540 | 474 | 802 |
| Total | 68 | 119 | 1,197 | 1,016 | 1,743 |

### Comparison between passive and active survey methods

Detection counts differed markedly between survey methods (Figure 3). The GLMM revealed a strong and significant effect of survey method on the number of *C*.*coturnix* detected. Active surveys using call playback detected significantly more individuals than passive surveys (β = 0.54 ± 0.04, *P* < 0.001) (Figure 4). On average, active detection increased expected counts by a factor of 1.72 (95% CI: 1.59–1.85) relative to passive surveys.

**FIGURE 3.**
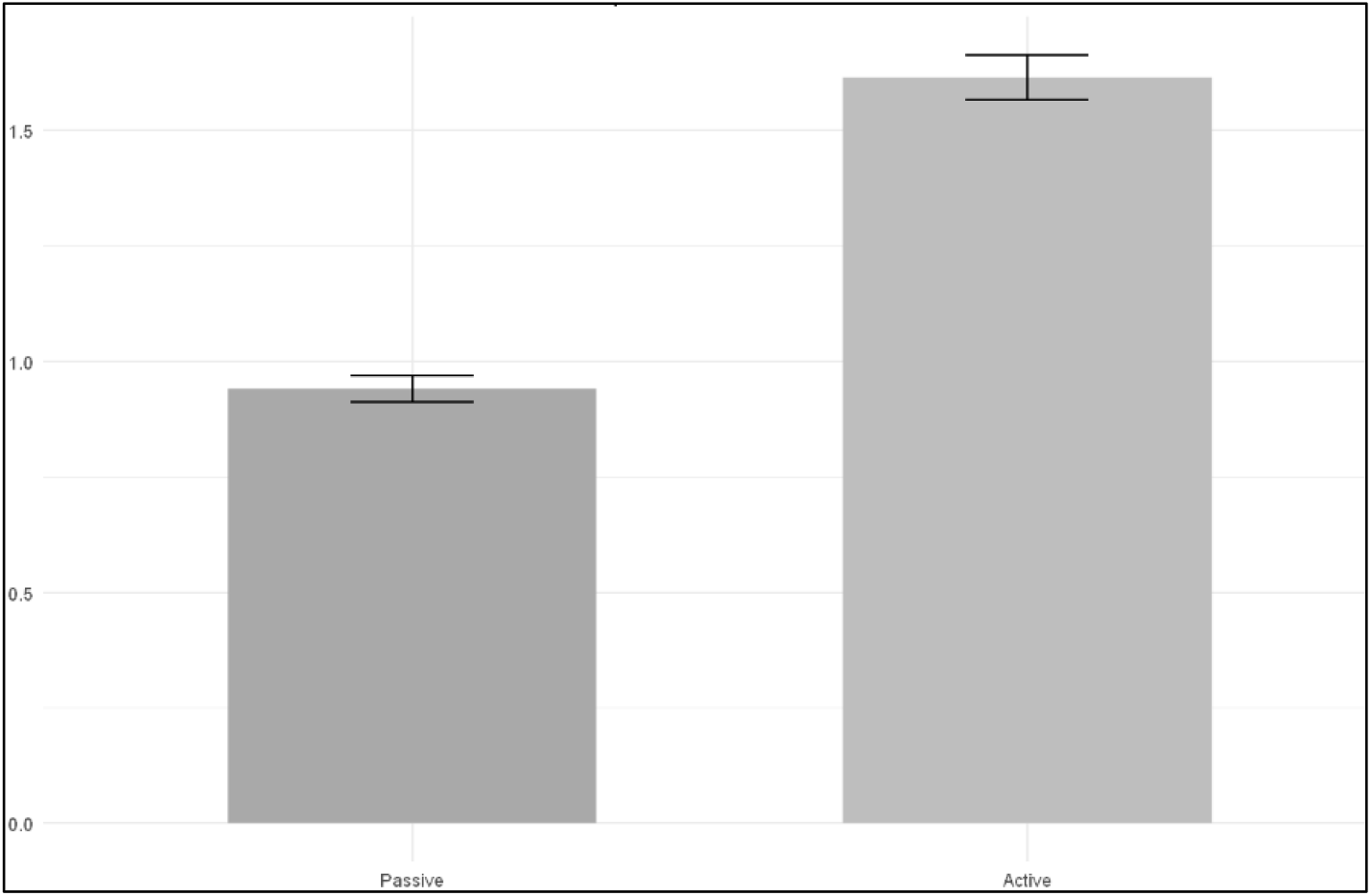
Increase in detection using passive (N_passive) vs. active (N_active) survey methods. Y-axis: Increase factor.

**FIGURE 4.**
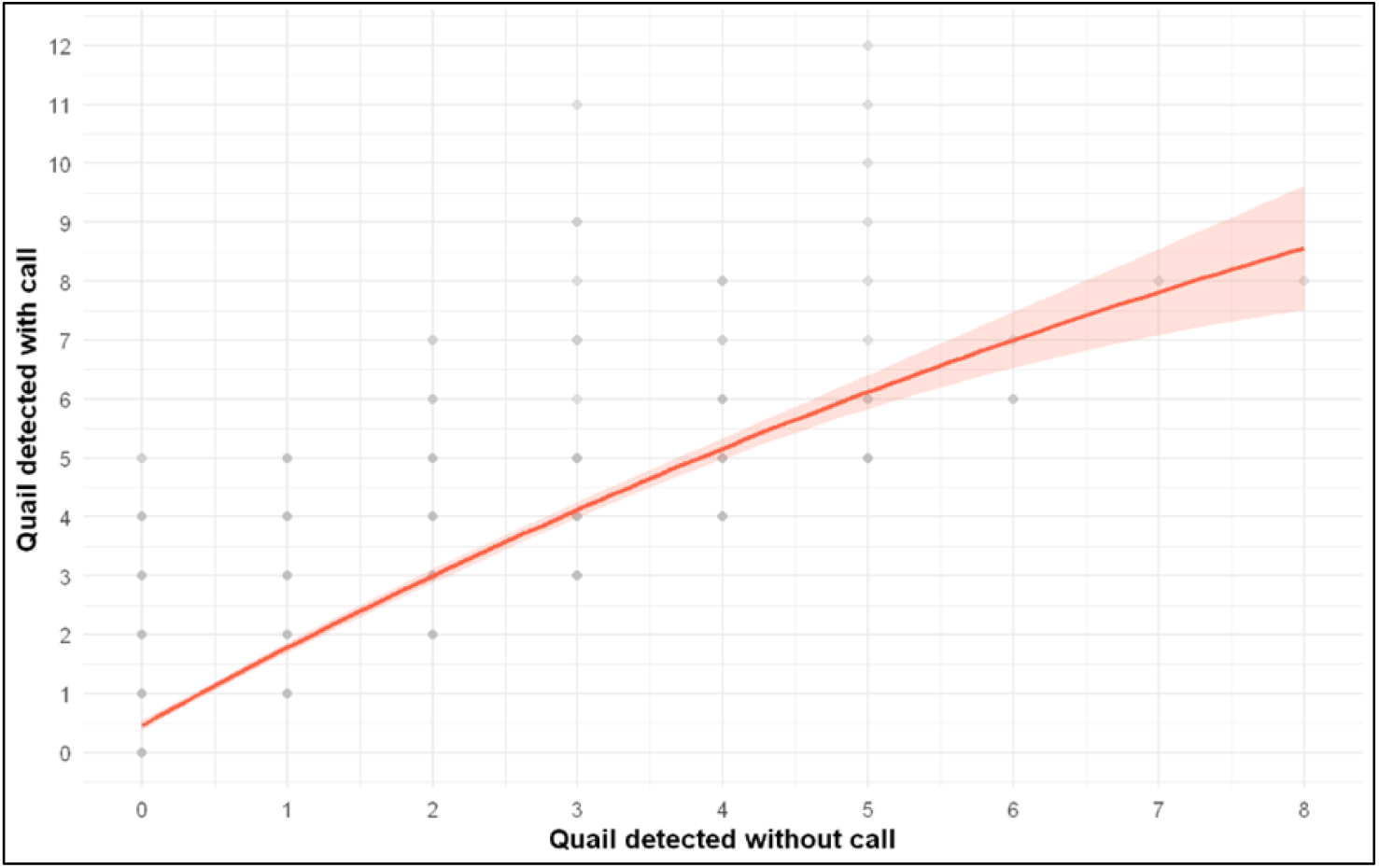
Relationship between passive (N_passive) and active (N_active) survey methods. The solid line represents the fitted GLMM, the shaded area shows the 95% confidence interval, and points represent observed values.

Model-based predictions confirmed this pattern, with higher mean predicted counts for active surveys across all survey transects and years. Observed versus predicted values showed good overall agreement, indicating adequate model fit. Marginal R^2^ values were low (R^2^m = 0.03–0.04), indicating that survey method alone explained only a small proportion of the variance in detection counts. In contrast, conditional R^2^ values were high (R^2^c = 0.61–0.74), showing that most of the explained variance arose from the hierarchical structure of the data, particularly differences among survey transects and sampling points. This result confirms strong non-independence among observations and highlights the limitations of analyses that ignore the nested sampling design.

### Relationship between passive and active detections

The increase in *C*.*coturnix* detections in response to playback (ΔN) showed a non-linear relationship with the number of individuals detected passively (GAM, edf = 2.26, *P* < 0.001, deviance explained = 5.6%, R^2^ = 4.4%) (Figure 5). ΔN increased with N_passive at low abundances, reaching a maximum of around 3–4 individuals per listening point, and at intermediate abundances, the increase stabilized (Figure 5).

**FIGURE 5.**
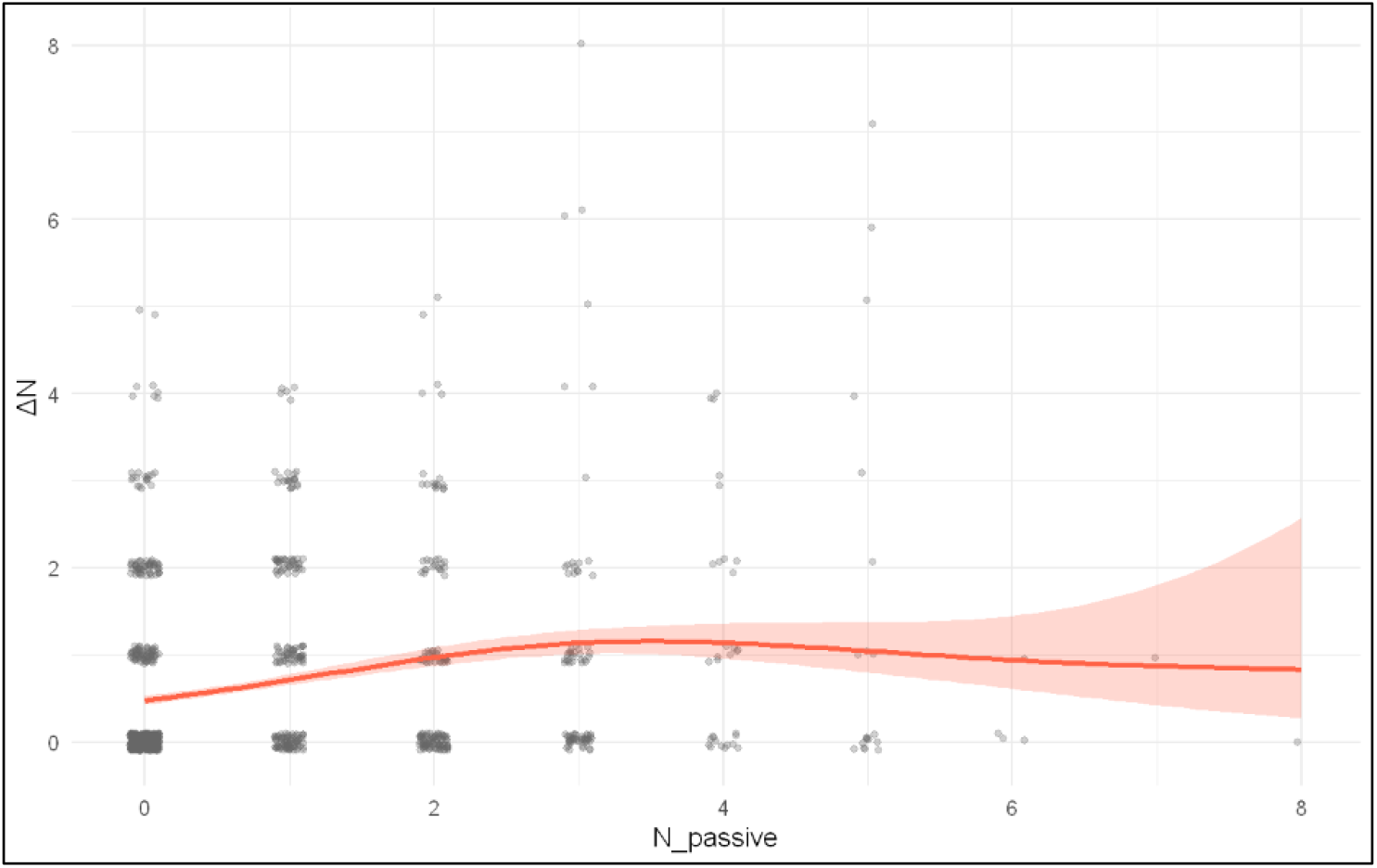
Relationship between increase in detections with claims and detections without claims. X-axis: Number of C.coturnix detected using passive method; Y-axis: Increase in C.coturnix detection using calls. Solid line: GAM estimate; shaded area: 95% confidence interval; points represent observed values.

## DISCUSSION

Breeding birds monitoring schemes are challenged when applied to species with cryptic behavior and variable spatial or temporal detectability. In this study, for the first time, we simultaneously compared the *C*.*coturnix* detectability using passive, generalist, multispecies surveys versus active and species-specific surveys employing female calls to elicit male responses. Active surveys consistently detected more individuals than passive ones and markedly improved detectability under low-abundance scenarios by recording individuals that would otherwise go undetected. This is particularly relevant in the current context, as assessments of the species’ conservation status in Europe rely on data from generalist and multispecies surveys (Arroyo et al. 2025), emphasizing that our results have important implications not only for the accurate monitoring of *C*.*coturnix* populations but also for their management and conservation.

### Comparison between passive and active survey methods

Active surveys using call playback consistently detected more individuals than passive surveys, with predicted counts on average 1.72 times higher. Previous research has similarly highlighted the limitations of passive surveys in optimal *C*.*coturnix* habitats. For example, Puigcerver et al. (2018), applying a protocol similar to ours in northeastern Spain, reported that passive methods detected only 10–30% of the individuals recorded by active surveys. Likewise, Sardà-Palomera et al. (2022) observed that active surveys increased *C*.*coturnix* detection from 21 to 36% relative to passive methods. Notably, in both studies, passive and active surveys were conducted on consecutive days, which may have affected results given the species’ high mobility (Munteanu and Maties 1974, Schleidt 1983, Rodríguez Teijeiro et al. 1992, Sardà-Palomera et al. 2011).

Evidence from other galliform species further supports the benefits of active methods: playback calls have been shown to increase detection probability in Black-breasted Wood-Quail (*Odontophorus leucolaemus*) (Hale, 2006) and help determine site occupancy in Northern Bobwhites (*Colinus virginianus*) (Duren et al. 2012), although these species differ from the *C*.*coturnix* in their ecology and movement patterns.

### Relationship between passive and active detections

Beyond the overall increase in detectability provided by active surveys, our analyses revealed a non-linear, density-dependent pattern. The increase in detections elicited by playback was largest at low passive abundances, reaching a maximum at around 3–4 individuals per listening point. At these abundances, active surveys substantially improve detection by recruiting individuals that would otherwise remain undetected. This pattern is consistent with the eco-ethological basis of *C*.*coturnix* vocal behaviour, in which male calling depends on social context and the presence and behavior of conspecifics (Pincemy and Guyomarc’h 2004), and many individuals may remain silent at low abundances unless stimulated. This pattern aligns with Sardà-Palomera et al. (2022), who also reported likely underestimation under low-abundance conditions. These results underscore the critical role of active, species-specific surveys in accurately detecting *C*.*coturnix*, particularly in low-abundance or spatially isolated populations, where individuals are less exposed to conspecific vocalizations and spontaneous calling is limited.

In addition, under low-abundance scenarios, which appear to predominate in Spain during the breeding season (Supplementary Material Table S2, Fundación Artemisan 2025), differences observed between the two methodologies may affect even basic presence–absence detection. In such contexts, monitoring based solely on passive methods may result in false absences, introducing additional uncertainty into assessments of species occurrence (Rodríguez-Teijeiro and Puigcerver 2022).

### Management implications

Our results demonstrate that active survey methods substantially enhance *C*.*coturnix* detectability compared to passive approaches, particularly under low-abundance conditions, which appear to predominate during the breeding season in Spain and are likely widespread across other European regions. These findings indicate that standard multispecies monitoring protocols may be insufficient to reliably capture *C*.*coturnix* presence and abundance, potentially introducing biases into population indices, trend estimates, and the detection of true demographic changes (Kéry and Schmidt 2008). Consequently, population trends for *C*.*coturnix* in Europe derived from passive monitoring and compiled by the PECBMS may be subject to substantial uncertainty (Arroyo et al. 2025).

Therefore, incorporating species-specific active methods, such as call playback, is critical to accurately assess *C*.*coturnix* presence and abundance, guide evidence-based management, and support adaptive harvest policies for a species of insecure status within the European Union (Cruz-Flores et al. 2025). Implementing active surveys across the *C*.*coturnix* European range will improve population modelling and enable robust conservation strategies, ensuring reliable assessments of population status and supporting the long-term sustainable conservation and responsible use of the species.

## Acknowledgements

We are indebted to game managers, hunters and volunteers for their help during surveys and landowners for allowing access to their farms. We are grateful to Santa Lambea and Diego A. Pulido for their help in field work and José D. Teijeiro for his comments and suggestions.

## Ethics statement

Permissions for land access were given by the landowners and the regional goverment issued permits for conducting surveys. As this was an observational study, no birds were captured.

## Author contribution

Conceptualization was carried out by E.L. and C.S.; project administration and funding acquisition by C.S.; supervision by R.C. and C.S.; investigation by E.L., R.C., J.A.T. and C.S.; coordination of field data collection by E.L., J.A.T. and M.B.; field data collection by J.R.; data curation by E.L. and I.N.; formal analysis by I.N.; visualization by E.L. and I.N.; and writing – review & editing by E.L., I.N., R.C., J.A.T. and C.S.

## LITERATURE CITED

Arroyo, B., M. Cruz-Flores, E. Šilarová, J. Škorpilová, M. Fernández-Tizón, and C. Carboneras (2025). Population update and model for Common Quail. https://circabc.europa.eu/ui/group/e21159fc-a026-4045-a47f-9ff1a319e1c5/library/4043c4ca-518a-4400-9ca1-2160558a4aa7/details

Bates, D., M. Maechler, B. Bolker, S. Walker, R.H.B. Christensen, H. Singmann, … and M.B. Bolker (2015). Package ‘lme4’. convergence, 12(1), 2.

Collins, S. A., and A. R. Goldsmith (1998). Individual and species diferences in quail calls (Coturnix c.japonica, C.c.coturnix and a Hybrid). Ethology 104:977–990.

Cruz-Flores, M., C. Carboneras, B. Rubio, M. Guillemain, J. Madsen, P. Defos du Rau, C. Francesiaz, and B. Arroyo (2025). Assessment of (un)sustainability of harvest. https://circabc.europa.eu/ui/group/e21159fc-a026-4045-a47f-9ff1a319e1c5/library/24f46784-5b20-4044-bab1-2df732f063f7/details

Derégnaucourt, S., and J. Guyomarc’h (2003). Mating Call Discrimination in Female European (Coturnix c . coturnix) and Japanese Quail (Coturnix c . japonica). Ethology 119:107–119.

Duren, K., J. Buler, W. Jones, and C. Williams (2012). Effects of Broadcasting Calls During Surveys to Estimate Density and Occupancy of Northern Bobwhite. Wildlife Society Bulletin 36:16–20.

Fundación Artemisan (2025). Monitorización y gestión de la codorniz común (Coturnix coturnix) en España. MEMORIA 2024–2025. https://fundacionartemisan.com/wp-content/uploads/2025/12/memoria-proyecto-coturnix-25.pdf

Gregory, R. D., A. Van Strien, P. Vorisek, A.W. Gmelig Meyling, D.G. Noble, R.P. Foppen, and D.W. Gibbons (2005). Developing indicators for European birds. Philosophical Transactions of the Royal Society B: Biological Sciences, 360(1454):269–288.

Guyomarc’h, H. (2003). Elements for a common quail (Coturnix c. coturnix) management plan. Game & Wildlife Science 20:1–92.

Guyomarc’h, J., A. Aupiais, and C. Guyomarc’h. (1998). Individual differences in the long-distance vocalizations used during pair bonding in European quail (Coturnix coturnix). Ethology Ecology & Evolution 10:333–346.

Hale, A. (2006). Group living in the black-breasted wood-quail and the use of playbacks as a survey technique. The Condor 108:107–119.

Holderried, P., H. Duschmalé, D. Günther, L. Isenberg, and J. Coppes (2025). Essential steps for establishing a large-scale passive acoustic monitoring for an elusive forest bird species: the Eurasian Woodcock (Scolopax rusticola). Ibis 167(2), 543–561.

Juan, M. (2012). Codorniz común-Coturnix coturnix. In: del Moral JC, Molina B, Bermejo A, Palomino D, editors. Atlas de las aves en invierno en España 2007–2010. Ministerio de Agricultura, Alimentación y Medio Ambiente–SEO/BirdLife, Madrid. Pages 116–117.

Kéry, M and B. Schmidt (2008). Imperfect detection and its consequences for monitoring in conservation. Community Ecology 9: 207–216.

Moussy, C., I.J. Burfield, P.J. Stephenson, A.F. Newton, S.H. Butchart, W.J. Sutherland, … and P.F. Donald (2022). A quantitative global review of species population monitoring. Conservation Biology 36(1), e13721.

Munteanu, D., and M. Maties (1974). The seasonal movements of the Quail (Coturnix coturnix L.) in Romania. Trav. Mus. Hist. Nat. ‘‘Grigore Antipa’’ 15: 365–380

Nichols, J. D., J.E. Hines, D.I. Mackenzie, M.E. Seamans, and R.J. Gutiérrez (2007). Occupancy estimation and modeling with multiple states and state uncertainty. Ecology 88(6):1395–1400.

Pincemy, G. and C. Guyomarc’h (2004). Inter-individual variability during morning choruses in Japanese quail males (Coturnix coturnix japonica). Comptes Rendus Biologies 1: 97–104

Puigcerver, M., C. Eraud, E. García-Galea, D. Roux, I. Jiménez Blasco, M. Sarasa, and J. Rodríguez-Teijeiro (2017). Common quail (Coturnix coturnix) in France and Spain: conflicting data or controversial census methodologies? In: Bro E, Guillemain M, editors. Proceedings 33 IUGB Congress and 13 Perdix. Montpelier, France.

Puigcerver, M., E. García-Galea, I. Jiménez-Blasco, S. Herrando, and J. D. Rodríguez-Teijeiro (2018). Mètodes generalistes passius de cens vs mètodes específics actius: el cas de la guatlla (Coturnix coturnix). Llibre de resums del 1r Congrés d’Ornitologia de les Terres de Parla Catalana. Barcelona.

Puigcerver, M., F. Sardà-Palomera, and J. D. Rodríguez-Teijeiro (2012). Determining population trends and conservation status of the common quail (Coturnix coturnix) in Western Europe. Animal Biodiversity and Conservation 35.2:343–352.

Puigcerver, M., F. Sardà-Palomera, and J. D. Rodríguez-Teijeiro. 2022. Codorniz común – Coturnix coturnix (Linnaeus, 1758). In: López P, Martín J, Casas F, editors. Enciclopedia Virtual de los Vertebrados Españoles. Museo Nacional de Ciencias Naturales, Madrid. https://www.vertebradosibericos.org/aves/cotcot.html

R Core Team (2024). R: A Language and Environment for Statistical Computing. R Foundation for Statistical Computing, Vienna, Austria. https://www.R-project.org/

Rodríguez-Teijeiro, J. D. and M. Puigcerver (2022). Codorniz común Coturnix coturnix. In: Molina B, Nebreda AR, del Moral JC, Muñoz AR, Seoane J, Real R, and J. Bustamante, editors. III Atlas de las aves en época de reproducción en España. SEO/BirdLife Madrid. https://atlasaves.seo.org/ave/codorniz-comun/

Rodríguez-Teijeiro, J. D., and M. Puigcerver (2020). Common Quail. In: Keller V, Herrando S, Voříšek P, Franch M, Kipson M, Milanesi P, Martí D, Anton M, Klvaňová A, Kalyakin MV, Foppen RP, editors. European Breeding Bird Atlas 2. Distribution, Abundance and Change. European Bird Census Council and Lynx Edicions, Barcelona.

Rodríguez-Teijeiro, J. D., F. Sardá-Palomera, I. Alves, Y. Bay, A. Beça, B. Blanchy, B. Borgogne, B. Bourgeon, P. Colaço, J. Gleize, A. Guerreiro, M. Maghnouj, C. Rieutort, D. Roux, and M. Puigcerver (2010). Monitoring and management of common quail Coturnix coturnix populations in their atlantic distribution area. Ardeola 57:135–144.

Rodríguez-Teijeiro, J., À. García, E. García-Galea, I. Jiménez-blasco, A. Torres-Riera, A. Barceló, M. Muñoz, F. J. Vidal, M. Puigcerver, and B. Seguí (2019). Dynamics of the population of Common quail males in the island of Majorca and comparison with the northeast peninsular populations. Mon. Soc. Hist. Nat. Balears 28:51–64.

Rodríguez Teijeiro, J. D., M. Puigcerver, and S. Gallego (1992). Mating strategy in the European quail (Coturnix c. coturnix) revealed by male population density and sex ratio in Catalonia (Spain) [mating system, polygyny spaced over time]. Gibier Faune Sauvage 9:377–386.

Sardà-Palomera, F., I. Jiménez-Blasco, M. Puigcerver, S. Herrando, and J. D. Rodríguez-Teijeiro (2022). ¿Cuántas codornices hay? Depende del método de censo. In: SEO-Birdlife, editors. Libro de Resúmenes del XXV Congreso Español de Ornitología. SEO/BirdLife, Madrid. pag 114.

Sardà-Palomera, F., Brotons, L., Villero, D., Sierdsema, H., Newson, S. E., and F. Jiguet (2012). Mapping from heterogeneous biodiversity monitoring data sources. Biodiversity and Conservation 21(11), 2927–2948.

Sardà-Palomera, F., M. Puigcerver, L. Brotons, and J. D. Rodríguez-Teijeiro (2012b). Modelling seasonal changes in the distribution of Common Quail Coturnix coturnix in farmland landscapes using remote sensing. Ibis 154:703–713.

Sardà-Palomera, F., Puigcerver, M. Vinyoles, D. & Sardà-Palomera, F. and J.D. Rodríguez-Teijeiro (2011). Exploring male and female preferences, male body condition, and pair bonds in the evolution of male sexual aggregation: the case of the Common Quail (Coturnix coturnix). Canadian Journal of Zoology 89: 325–333.

Schleidt, W.M. (1983). Spatial and temporal Pattern of Calling Sites in Coturnix Quails. Nat. Gro. Soc. Res. Rep. 15: 573–576.

