## Supplementary 1 for "Playback calls help to increase the detectability of *Coturnix coturnix* (Common quail), a cryptic and widespread galliform"

### Supplementary material Table S1. Paper-based protocol.

**QUAIL SURVEY DATA WITH FEMALE CALL PLAYBACK** **Province:**___________ **Locality:** ___________ **Survey ID**:__________

**Date:** _________ **Observer:**_________________________ **Start time:**_______ **End time:**_______ **UTM Grid:**
**Temperature:** ☐ cold ☐ mild ☐ hot **Sky:** ☐ cloudy ☐ partly cloudy ☐ clear **Wind:** ☐ no wind ☐ breeze ☐ windy ☐ strong wind
**Ground:** ☐ dry ☐ wet ☐ waterlogged **Precipitation**: ☐ storm ☐ hail ☐ drizzle ☐ fog ☐ none

| Listening point | Time | QUAILS DETECTED | CROP | | | | OBSERVATIONS |
|---|---|---|---|---|---|---|---|
| | | | Type | Height (cm) | Color | Ripening | |
| 1 | | | | | | | |
| 1c | | | | | | | |
| 2 | | | | | | | |
| 2c | | | | | | | |
| 3 | | | | | | | |
| 3c | | | | | | | |
| 4 | | | | | | | |
| 4c | | | | | | | |
| 5 | | | | | | | |
| 5c | | | | | | | |
| 6 | | | | | | | |
| 6c | | | | | | | |
| 7 | | | | | | | |
| 7c | | | | | | | |

| |
|---|
| 8 |
| 8c |
| 9 |
| 9c |
| 10 |
| 10c |

**1,2,3,4….10: passive listening period ; 1c,2c,3c….10c: active listening period**

**Crop: Type:** B barley, W wheat, O oats, P pea, V vetch, S sunflower … ; **Height** (cm); **Color**, G (green) G2 (golden); **Ripening:** Yes / No
