## Supplementary 2 for "Playback calls help to increase the detectability of *Coturnix coturnix* (Common quail), a cryptic and widespread galliform"

### Supplementary material Table S2. Survey transects in the Hunter's Watch

The Hunter's Watch (in Spanish *Observatorio Cinegético*, hereafter, HW) is a voluntary based project in which hunters and game managers are invited to conduct wildlife monitoring, primarily on game species. It consists of a mobile application to record field observations and a web platform where the information is received and analysed afterwards. This project aims to address the distribution, abundance, and in the medium-to-long term, population dynamics and trends of game species.

#### Methodology

Monitoring is conducted through surveys, mainly transects by foot or vehicle. In the case of the *Coturnix coturnix* common quail, these sruveys are part of the summer migratory bird campaign, alongside the *Streptopelia turtur* European turtle dove. For this campaign, collaborators are recommended to select a route (4–6 km in length) that covers partially favorable areas during breeding periods for the target species, recording all detected individuals (both seen and heard). The technical staff of Artemisan provides assistance to volunteers regarding route selection and the use of the mobile application. In addition to volunteers, some professional field researchers also carry out this work.

In this study, summer migratory bird censuses were conducted between April and August from 2021 to 2025. Both the survey route and the observations were recorded using the CensData app, available for Android and iOS. After data exploration, surveys shorter than 3 km were discarded from further analyses, as they were considered not representative of the hunting ground or the 10×10 km UTM grid in which conducted. The total number of observations was divided by the kilometers travelled to obtain the Kilometric Abundance Index (KAI).

#### Results

Between April and August from 2021 to 2025, 1,901 linear transects of summer migratory birds were conducted in 696 different 10×10 km UTM grids (112 in 2021, 231 in 2022, 262 in 2023, 315 in 2024, and 423 in 2025), detecting 5,902 quails (Figure S1). The average KAI for quail was 0.49 quails/km, with very wide standard deviations. In 59.92% of the sampled grids, no quail were detected.

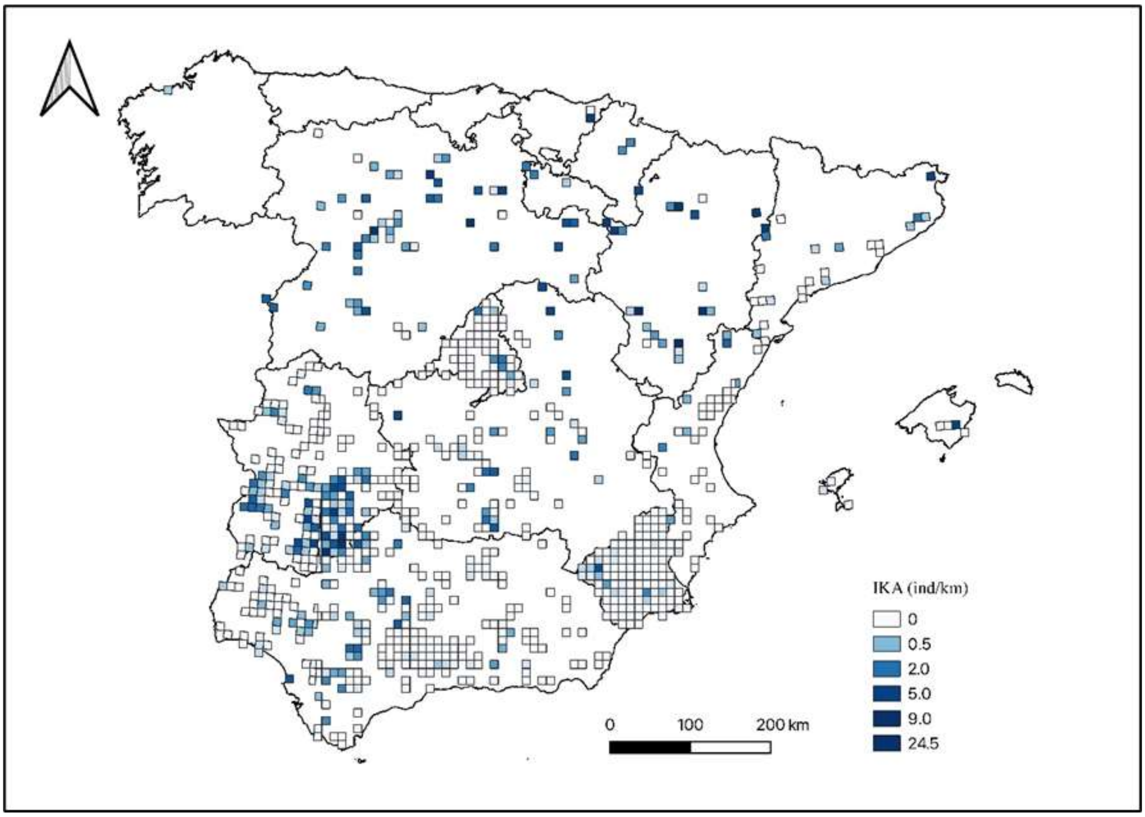


**Figure S1.** Map showing the 696 10×10 km UTM grids where summer migratory bird surveys were conducted from 2021 to 2025 through the HW, colored according to the Kilometric Abundance Index (IKA) of quail obtained.
